# Modelling molecular differences in the innate immune system responses of chickens and ducks to highly pathogenic avian influenza virus

**DOI:** 10.1101/2024.07.26.605270

**Authors:** Tamsin Wood, Gary An, Clare E. Bryant, Brian J. Ferguson

## Abstract

Highly pathogenic avian influenza virus (HPAIV) presents a global threat to chicken livestock; chickens infected by HPAIV tend to show severe symptoms and high mortality rates. In 2022, the largest recorded outbreak of HPAIV in Europe resulted in millions of chickens being culled in the UK alone to try to prevent further spread. Unlike chickens, mallard ducks show reduced symptom severity and lower mortality rates to HPAIV infection. Research into the immune system responses of these two species shows they differ in their molecular outputs: chickens produce a pro-inflammatory response; mallards produce an anti-viral response. These differences in immune responses are thought to be in part due to chickens missing pattern recognition receptor retinoic acid-inducible gene-I (RIG-I). This project aimed to model the innate immune systems of chickens and mallard ducks to an abstracted molecular level. A literature search was conducted, and the immune systems were modelled in NetLogo as an avian innate immune response agent-based model (AIIRABM). The AIIRABM enabled examination of the relative importance of molecular differences between the chicken and mallard duck innate immune systems and produced similar differences in chicken and mallard duck molecular outputs to those observed *in vitro* and *in vivo*. Simulation experiments with the AIIRABM supported the molecular difference RIG-I as key in causing the differences in the chicken and mallard duck innate immune responses to HPAIV. The AIIRABM will be used in further research on the chicken and mallard duck immune responses to HPAIV as the baseline in an iterative modelling cycle.

## Introduction

Avian influenza virus (AIV) refers to any Influenza A strain with birds as the natural reservoir (Causey & Edwards, 2008). Some AIV strains are zoonotic (Kalthoff *et al*, 2010) and as such the World Health Organisation acknowledges the potential for AIV to induce a pandemic within the human population (Thomas & Noppenberger, 2007). However, so far, no AIV outbreaks in humans have shown sustained human-to-human transmission (Zhou *et al*, 2018). AIV is panzootic in both domestic and wild bird populations (Ramey *et al*, 2022). In 2022, Europe saw its largest AIV epidemic yet with thousands of outbreaks reported in both domestic and wild avian populations (EFSA, ECDPC, EURLAI *et al*, 2023). The UK alone culled over four million farmed chickens to attempt to minimise the epidemic’s spread (Haider *et al*, 2023) and AIV devastated some wild bird populations including UK seabirds (Tremlett *et al*, 2024). In 2024 AIV spread further into multiple mammalian species, including dairy herds in the USA (Nguyen *et al*, 2024), and has broad lethality in numerous populations of wild mammals, especially carnivores where transmission is assumed to be via ingestion of infected birds (Plaza *et al*, 2024). Studying the spread and effect of AIV is therefore highly important to minimise its negative effects on wild bird populations and economic losses in poultry agriculture.

Not all AIV strains cause high mortality in birds due to differences in viral proteins (Pantin-Jackwood & Swayne, 2009). Consequently, every AIV strain is categorised as either highly pathogenic AIV (HPAIV) or low pathogenic AIV (LPAIV) based on its severity of symptoms and degree of lethality in chickens (*Gallus gallus domesticus*) (Swayne, 2007). With chickens as the major poultry livestock worldwide (Scanes, 2018), understanding the effect of HPAIV in this species is essential to minimising food losses and potential impacts on global food security. Unlike chickens, mallard ducks (*Anas platyrhynchos*) often develop less severe symptoms and display lower mortality rates with HPAIV infection (Evseev & Magor, 2019; Uchida *et al*, 2019; Burggraaf *et al*, 2014). (From now on, unless otherwise stated, “duck” refers to “mallard duck”). Frequently chickens experimentally infected with HPAIV die more rapidly and at a higher mortality rate compared to ducks (Alexander *et al*, 1986; Burggraaf *et al*, 2014; Jeong *et al*, 2009). This difference in species is not quite so simple as HPAIV infectivity differs between chicken and duck breeds (Matsuu *et al*, 2016; Sánchez-González *et al*, 2020; Pantin-Jackwood & Suarez, 2013), but understanding why ducks generally display reduced infection severity could inspire modifications to the chicken immune system to reduce the negative effect of HPAIV.

How ducks better respond to HPAIV compared to chickens is an area of active research. With both natural and experimental infection, HPAIV shows distinct virus distributions in chicken and duck organ tissues (Vreman *et al*, 2022). Furthermore, chicken and duck immune responses to HPAIV vary with tissue type (Cornelissen *et al*, 2013; Watanabe *et al*, 2011). Perhaps these differential tissue-level responses to infection result in the difference in HPAIV severity between chickens and ducks. At the cellular level additional differences in avian responses to HPAIV infection are observed. In particular, HPAIV readily replicates in chicken endothelial cells and induces high production levels of pro-inflammatory cytokines but the same phenomenon is not observed in ducks (Vreman *et al*, 2022; de Bruin *et al*, 2022; Tong *et al*, 2021). The more severe infection of endothelial cells in chickens compared to ducks could enable greater spread of AIV to other tissues and therefore also account for the species differences in HPAIV infection.

As well as tissue and cell type variation, the molecular products of the general innate immune response to HPAIV differ between chickens and ducks. With AIV infection virus production is often initially comparable between chicken and duck tissues, but by 1 day post infection (d.p.i.) chicken cells can be producing four times as many virions as duck cells (Al-Mubarak *et al*, 2015). Furthermore, pro-inflammatory cytokines production is greater in chickens than ducks (Kuchipudi *et al*, 2014; de Bruin *et al*, 2022) and is thought to in part account for the severity of HPAIV symptoms (Kuribayashi *et al*, 2013). The type-1 interferon (IFN) response also often differs between chickens and ducks with AIV infection (Cornelissen *et al*, 2013; El-Shall *et al*, 2023; Liang *et al*, 2011) and as type-1 IFNs stimulate an anti-viral cell state (Evseev & Magor, 2019; Kuchipudi *et al*, 2014) it is possible variation in this response could further account for the difference in HPAIV severity between chickens and ducks.

Key molecular differences between the chicken and duck innate immune systems could be responsible for most of the variation observed in their molecular outputs with HPAIV infection. For example, absence of retinoic acid-inducible gene-I (RIG-I) from chicken pattern recognition receptors is thought to be key to ducks’ greater HPAIV immunity; RIG-I affects the production of many key immune response molecules (Karpala *et al*, 2011; Barber *et al*, 2010). Transfection of duck RIG-I (duRIG-I) into chickens has reduced HPAIV replication and altered the production of various key immune response molecular outputs (Barber *et al*, 2010, 2013; preprint: Sid *et al*, 2023).

An avian innate immune response agent-based model (AIIRABM) was created to compare the chicken and duck systems. The AIIRABM generally showed similar differences in chicken and duck molecular outputs as those observed experimentally. Additionally, *in silico* testing with the AIIRABM confirmed the effect of transfecting duRIG-I into chickens; the duRIG-I chicken immune response more closely resembled the duck response. The AIIRABM created in this project will be used as the baseline in a future iterative modelling cycle, aiding further research on the molecular differences in the chicken and duck innate immune responses to HPAIV.

## Materials and Methods

### Aggregation and abstraction of the scientific literature

The AIIRABM focused on epithelial cells and M1 macrophages within the avian innate immune response due to their critical nature in sensing AIV. To produce the abstracted agent-based model of the chicken and duck innate immune systems a scientific literature search was conducted on the known elements of the avian innate immune response to HPAIV and any known molecular differences between the two systems. Where research was limited on a pathway or element, scientific knowledge of the mouse innate immune system was cautiously used instead. This was particularly relevant when determining the key molecular outputs of cell types; limited to no research has been conducted on the key molecular outputs of avian innate immune cells. The scientific literature search produced an outline of the avian innate immune response to a molecular level within epithelial cells and M1 macrophages. Two key molecular differences were identified between the chicken and duck systems: (1) the absence of RIG-I and its coreceptor RNF135 in chickens (preprint: Sid *et al*, 2023); (2) greater NLRP expression in chickens than in ducks (Campbell & Magor, 2020).

The avian innate immune systems were abstracted based off the desired utility of the AIIRABM. The AIIRABM was built to predict the system effects of key known molecular differences between chickens and ducks, to determine whether they could reproduce the differences in chicken/duck innate immune responses observed with *in vitro* cell culture and *in vivo* experiments. Furthermore, the AIIRABM was produced as the baseline model of an iterative modelling cycle; the AIIRABM will inform a series of *in vitro* cell culture experiments that will either concur or contradict the model, inspiring another iteration of the AIIRABM and so on. Based off these desired functions the avian innate immune systems were simplified into the main molecular pathways of both cell types and only key molecular elements of each pathway were modelled. For example, the pathway from pattern recognition receptor MDA5 to type-1 IFNs secretion (including molecular elements MAVS, TBK1, IKKε, etc.) was abstracted to just MDA5 activation, IRF7 activation, and type-1 IFNs production and secretion. From this abstraction a cell state diagram was produced for both cell types (Fig.1).

**Figure 1:**
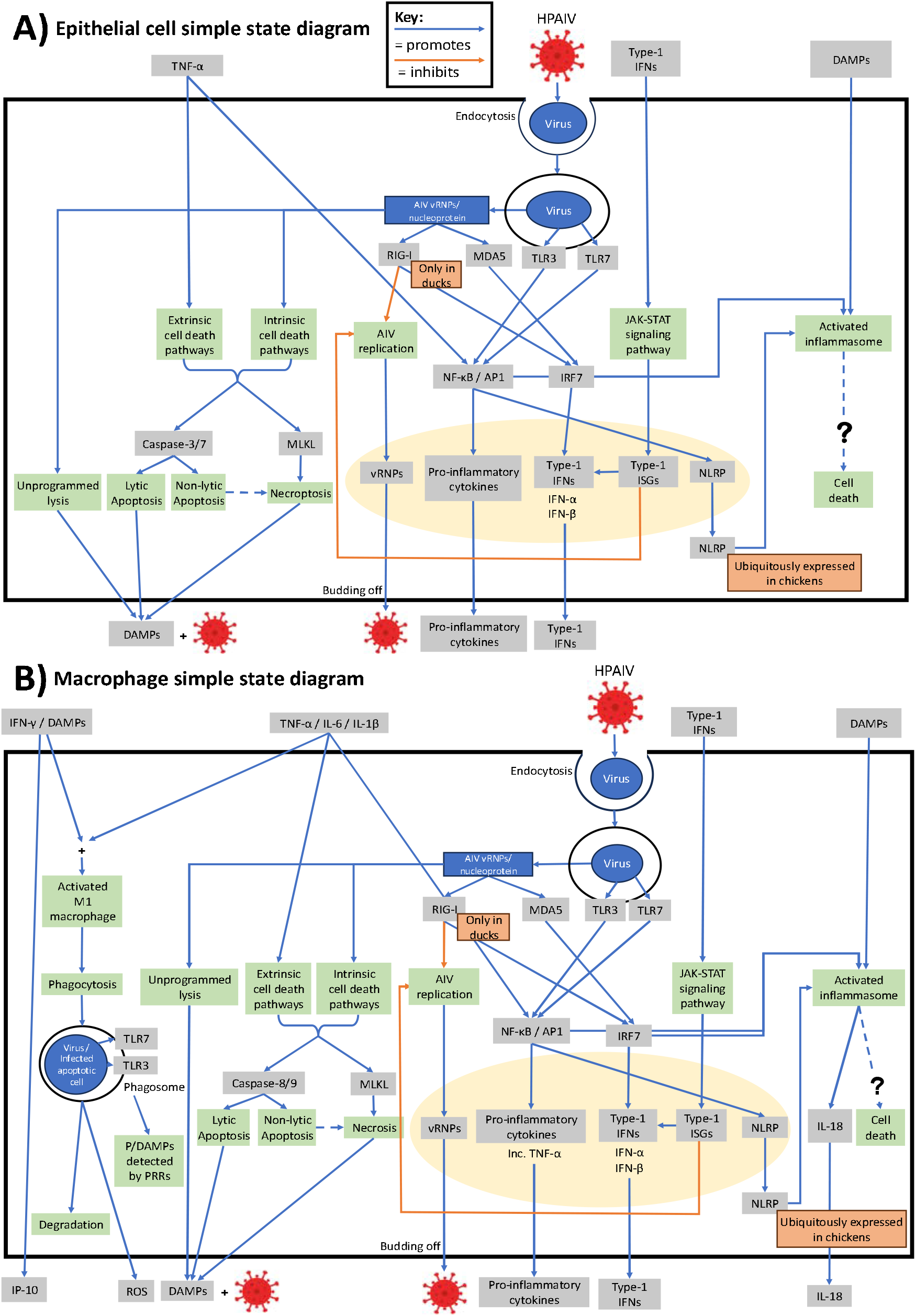
Cell state diagrams of the abstracted innate immune responses to HPAIV in avian epithelial cells (**A**) and macrophages (**B**). Arrows between molecular elements represent either activating (blue) or inhibitory (orange) interactions. The two key molecular differences between the chicken and duck systems are shown in the orange labels. The yellow oval represents the cell nucleus.

### The NetLogo model

Agent-based modelling is a good technique for representing immune systems as it allows for complex, non-linear, multi-agent interactions and can produce unexpected model outputs inspiring novel hypotheses (Chiacchio *et al*, 2014). The AIIRABM was encoded in NetLogo (Wilensky, 1999), a free open-source programming language and IDE designed specifically for users to conduct agent-based modelling relatively easily. The NetLogo files for the original AIIRABM and its RIG-I MDA5 variants are available upon request.

The AIIRABM consists of a 33 × 33 grid of “patches” (NetLogo terminology) with a fixed epithelial cell initially on each patch and mobile macrophages initially randomly placed (Fig.2). This model topology aims to mimic *in vitro* cell culture experiments. Epithelial cells and macrophages are “turtles”, distinguished by “breed”. The AIIRABM updates with each “tick”. With calibration 360 ticks are equivalent to 1 d.p.i. All AIIRABM simulations terminate after 2 d.p.i. to match cell culture experimental timeframes.

**Figure 2:**
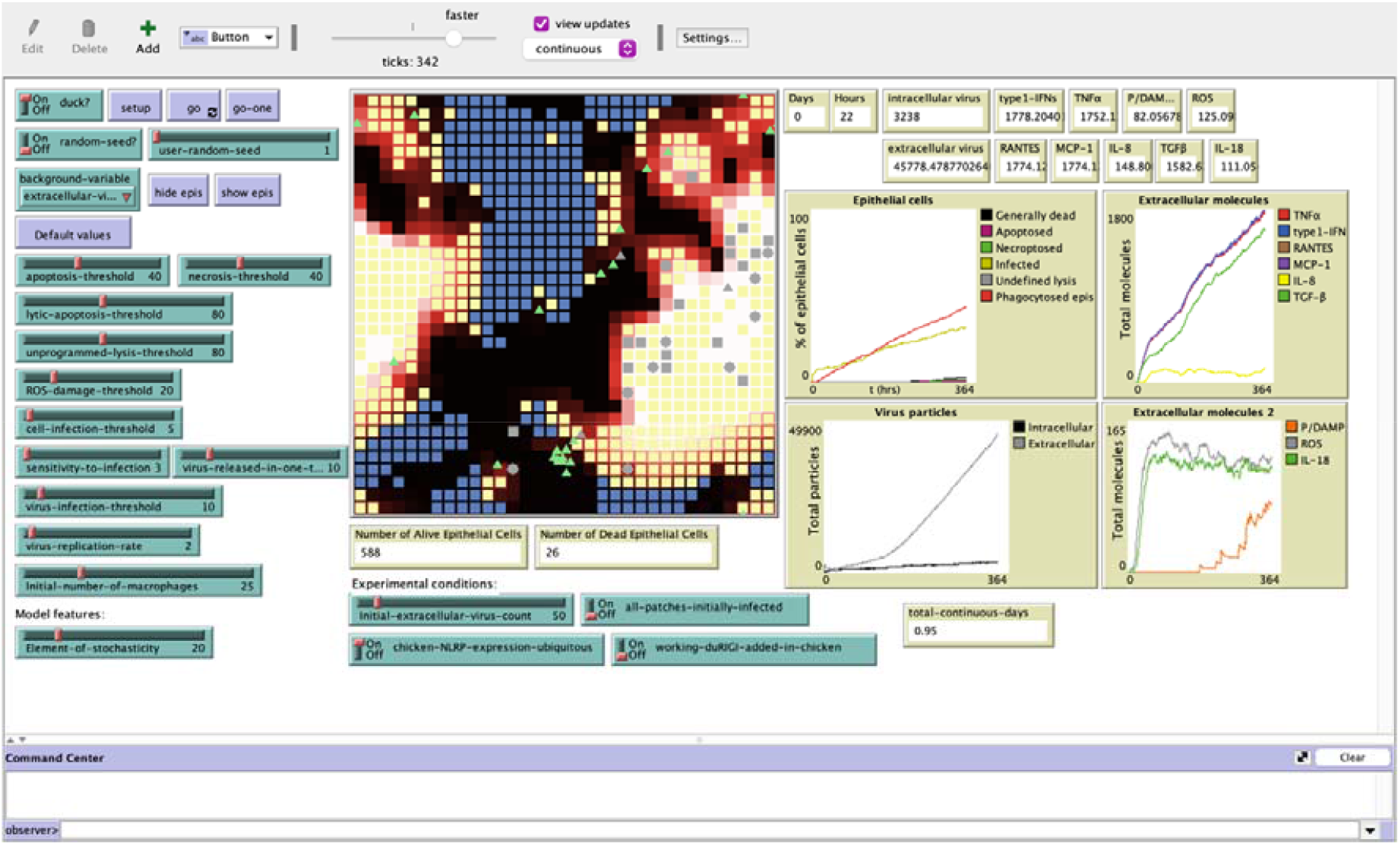
The NetLogo graphical user interface (GUI) with AIIRABM mid-simulation. A 33 × 33 grid of “patches”, each with a fixed epithelial cell (squares) and mobile macrophages initially randomly placed in the grid (triangles). An epithelial cell is blue when uninfected by HPAIV, yellow when infected and grey when dead. An epithelial cell is petal-shaped when apoptotic. A patch is blank when its epithelial cell has been phagocytosed. A macrophage is purple when uninfected by HPAIV, green when infected and grey when dead. The degree of background red represents the amount of extracellular virus incident on a patch; a white background represents the greatest amount of extracellular virus incident. The switches and sliders on the GUI enable the user to easily change the set of parameters used in the model.

Each element of the abstracted molecular pathways of epithelial cells and macrophages is represented in the AIIRABM by either a “patch variable” or “turtle variable” (Table.1). Patch variables represent the amount of extracellular element on a given patch such as total extracellular virions, type-1 IFNs, etc. Turtle variables represent the intracellular molecular elements of molecular pathways. The value of a molecular element’s turtle variable is dependent upon the turtle variable(s) of the molecular element(s) one above in the pathway. With each tick all patch and turtle variables are updated. Within a tick the last element in a molecular pathway is the first to have its turtle variable updated to ensure that the whole molecular pathway does not always run in one tick.

**Table 1:**
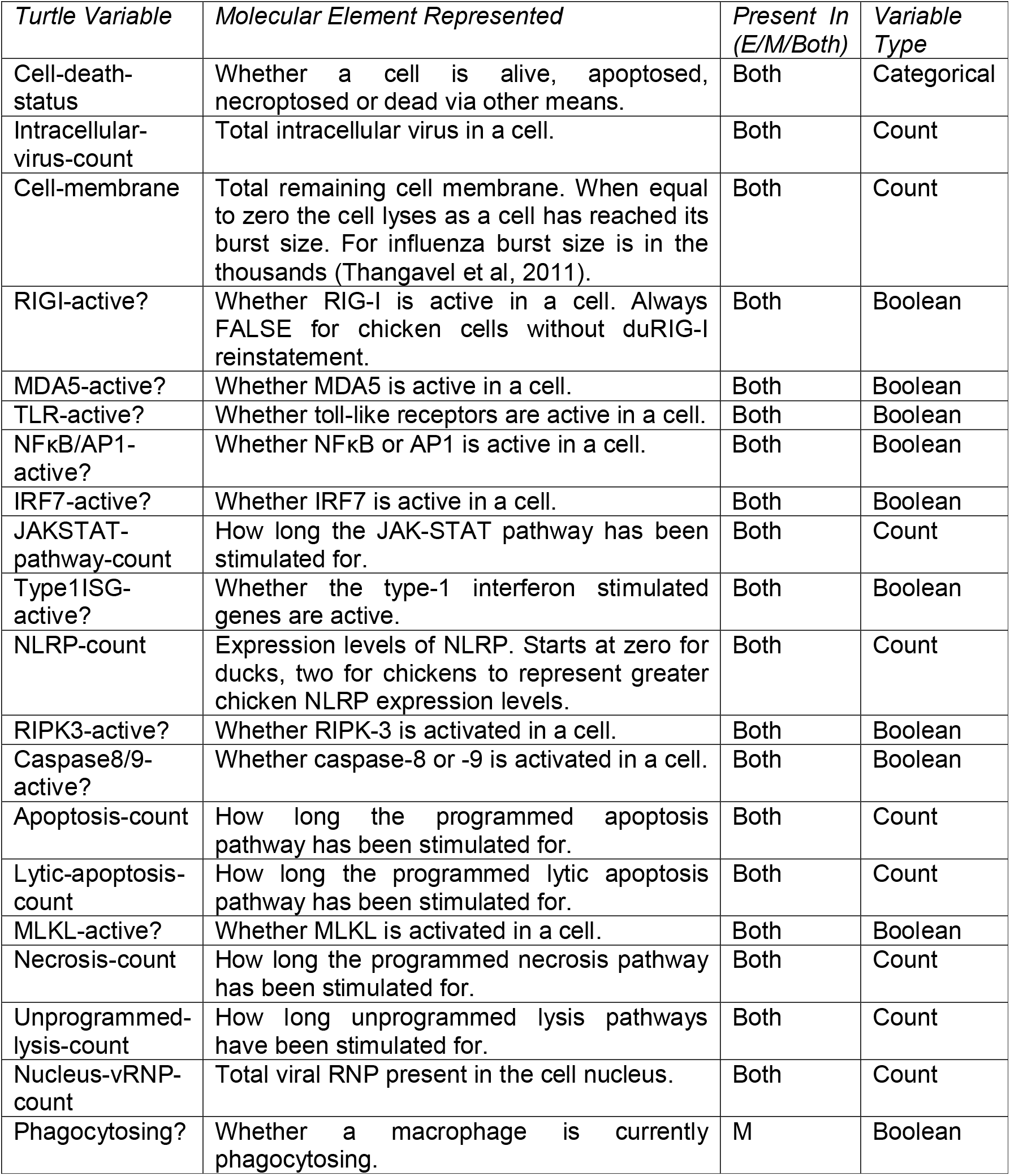
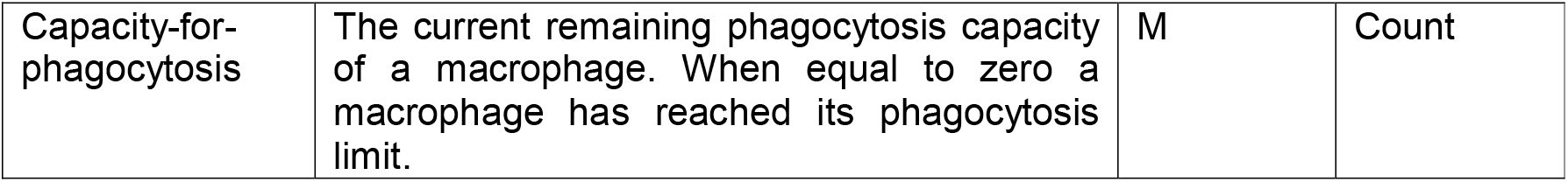
Turtle variables in the AIIRABM and the molecular elements they represent. For the ‘Present In’ column, ‘E’ stands for ‘epithelial cells’ and ‘M’ stands for ‘macrophages’.

| *Turtle Variable* | *Molecular Element Represented* | *Present In (E/M/Both)* | *Variable Type* |
| --- | --- | --- | --- |
| Cell-death-status | Whether a cell is alive, apoptosed, necroptosed or dead via other means. | Both | Categorical |
| Intracellular-virus-count | Total intracellular virus in a cell. | Both | Count |
| Cell-membrane | Total remaining cell membrane. When equal to zero the cell lyses as a cell has reached its burst size. For influenza burst size is in the thousands (Thangavel et al, 2011). | Both | Count |
| RIGI-active? | Whether RIG-I is active in a cell. Always FALSE for chicken cells without duRIG-I reinstatement. | Both | Boolean |
| MDA5-active? | Whether MDA5 is active in a cell. | Both | Boolean |
| TLR-active? | Whether toll-like receptors are active in a cell. | Both | Boolean |
| NFkB/AP1-active? | Whether NFkB or AP1 is active in a cell. | Both | Boolean |
| IRF7-active? | Whether IRF7 is active in a cell. | Both | Boolean |
| JAKSTAT-pathway-count | How long the JAK-STAT pathway has been stimulated for. | Both | Count |
| Type1ISG-active? | Whether the type-1 interferon stimulated genes are active. | Both | Boolean |
| NLRP-count | Expression levels of NLRP. Starts at zero for ducks, two for chickens to represent greater chicken NLRP expression levels. | Both | Count |
| RIPK3-active? | Whether RIPK-3 is activated in a cell. | Both | Boolean |
| Caspase8/9-active? | Whether caspase-8 or -9 is activated in a cell. | Both | Boolean |
| Apoptosis-count | How long the programmed apoptosis pathway has been stimulated for. | Both | Count |
| Lytic-apoptosis-count | How long the programmed lytic apoptosis pathway has been stimulated for. | Both | Count |
| MLKL-active? | Whether MLKL is activated in a cell. | Both | Boolean |
| Necrosis-count | How long the programmed necrosis pathway has been stimulated for. | Both | Count |
| Unprogrammed-lysis-count | How long unprogrammed lysis pathways have been stimulated for. | Both | Count |
| Nucleus-vRNP-count | Total viral RNP present in the cell nucleus. | Both | Count |
| Phagocytosing? | Whether a macrophage is currently phagocytosing. | M | Boolean |
| Capacity-for-phagocytosis | The current remaining phagocytosis capacity of a macrophage. When equal to zero a macrophage has reached its phagocytosis limit. | M | Count |

AIIRABM simulations were run in the NetLogo BehaviorSpace. 10 replications were conducted per simulation setup across random seeds 1-10.

### Statistical analysis

All statistical analyses were conducted using base R functions (R Core Team, 2023). ANOVA normality assumptions were tested with the Shapiro-Wilk test.

## Results

### Calibration and comparison of the chicken and duck AIIRABMs

The AIIRABM was calibrated to avian AIV *in vitro* cell culture experiments and produced biologically sensible behaviours: extracellular virus count increased to a peak and subsequently decreased during the 2-day simulation period (de Bruin *et al*, 2022); type-1 IFNs were upregulated by 1 d.p.i. (Kuchipudi *et al*, 2014); pro-inflammatory cytokine production increased and decreased along with the viral dynamics (Cornelissen *et al*, 2013).

The AIIRABM was run across two days to mimic AIV *in vitro* cell culture experiments. Both chicken and duck AIIRABMs were simulated at low initial inoculum (initial virus count = 50) with 10 stochastic replicates. Total extracellular virus in both AIIRABMs initially increased, peaked, and subsequently decreased across the 2-day simulation period (Fig.3). Total extracellular virus peak intensity in the chicken AIIRABM was more than double the peak intensity in the duck AIIRABM (one-way ANOVA, p < 0.05). Furthermore, total extracellular virus peaked earlier in the chicken system than the duck (one-way ANOVA, p < 0.05); chicken extracellular virus peaked before 1 d.p.i. unlike the duck peak. The duck total extracellular virus curve did not return to zero arbitrary units by 2 d.p.i. unlike the chicken curve. This is because not all epithelial cells died in each duck stochastic replicate by 2 d.p.i. whereas nearly all epithelial cells died by 2 d.p.i in the chicken AIIRABM.

**Figure 3:**
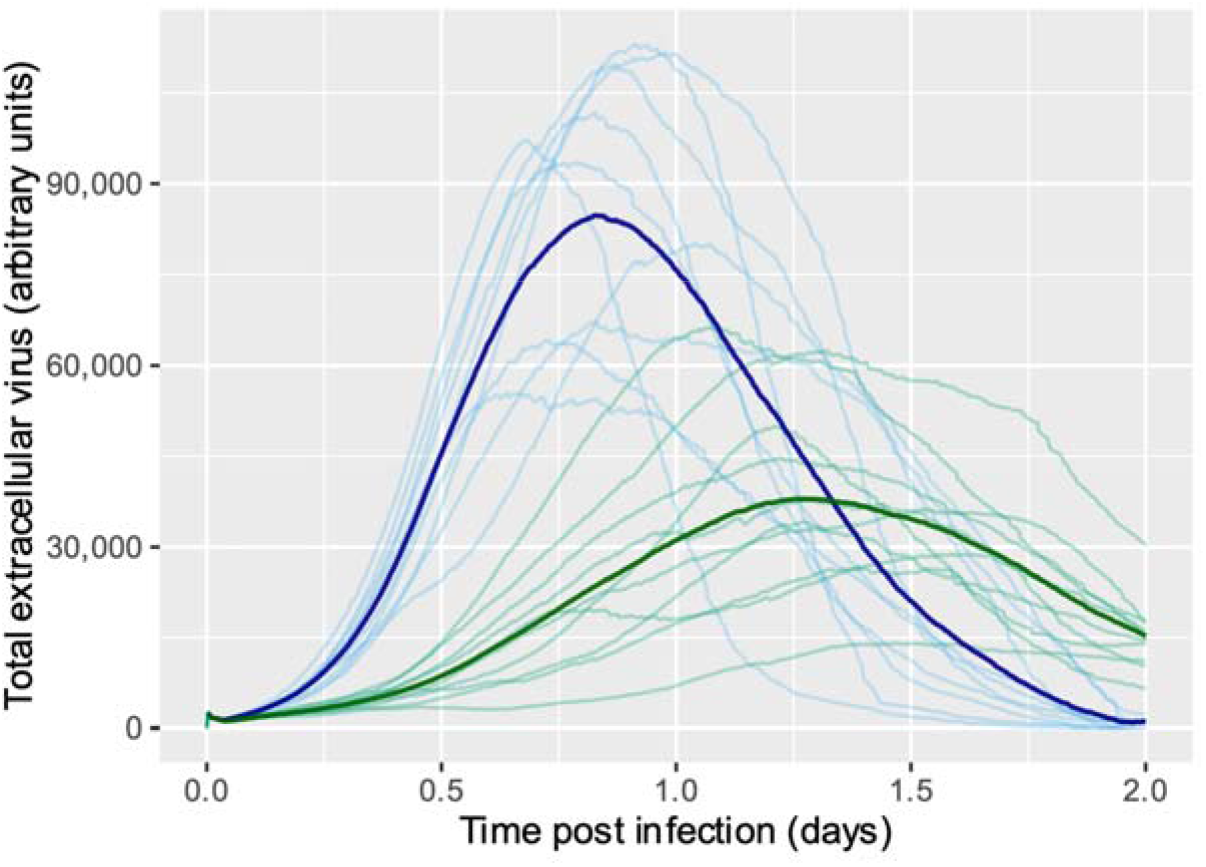
The AIIRABM calibrated, simulated across two days and at low initial inoculum (initial virus count = 50). Trajectories of total extracellular virions across time in 10 stochastic replicates and their average are plotted for both the chicken (blue) and duck (green) innate immune systems.

TNF-α production in both chicken and duck AIIRABMs initially rapidly increased, peaked and subsequently decreased (Fig.4A). Chicken total TNF-α peak intensity was nearly double duck peak intensity (one-way ANOVA, p < 0.05). Additionally, the chicken TNF-α peak occurred much earlier than the duck peak (Kruskal-Wallis, p < 0.05). Total type-1 IFNs in both chicken and duck AIIRABMs initially rapidly increased, then increased to a peak, before subsequently decreasing (Fig.4B). Total type-1 IFNs peak intensity was much greater in the duck than in the chicken AIIRABM (Kruskal-Wallis, p < 0.05). Additionally, the duck type-1 IFN response was maintained over a greater period than the chicken response; total type-1 IFNs in the chicken AIIRABM peaked and started declining by 1 d.p.i. whereas duck type-1 IFNs on average peaked after 1 d.p.i.

**Figure 4:**
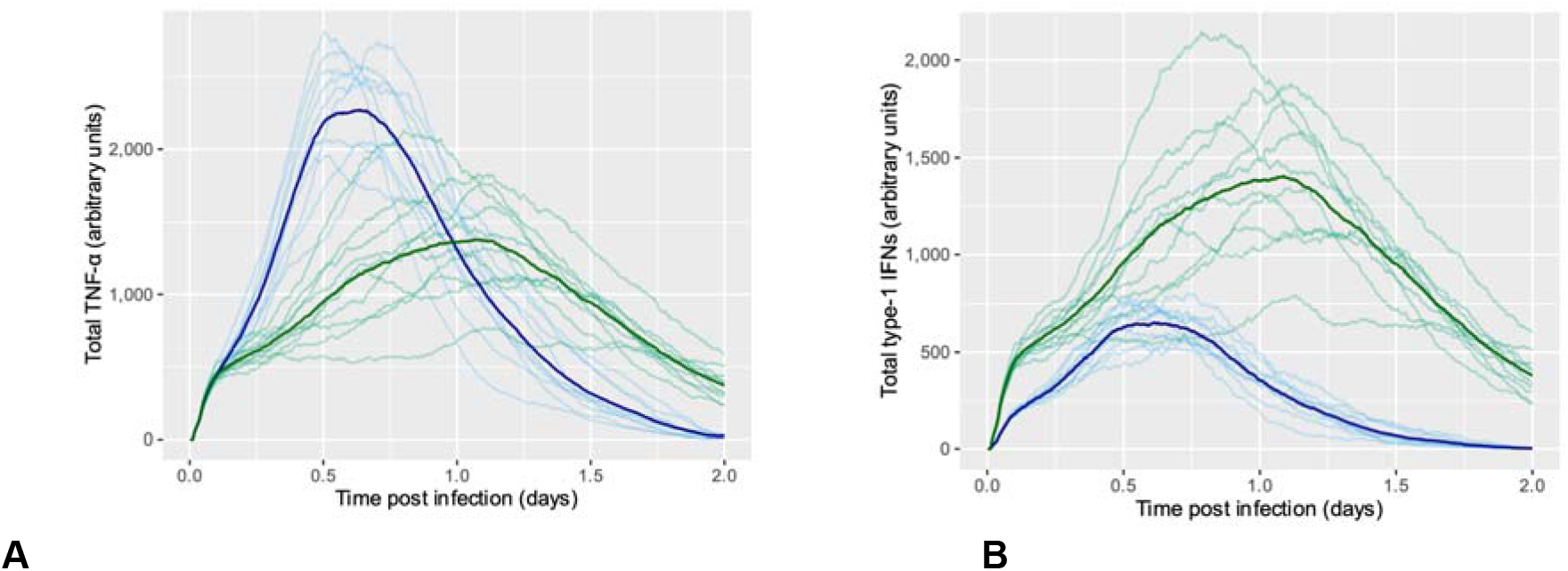
The AIIRABM calibrated, simulated across two days and at low initial inoculum (initial virus count = 50). Trajectories of total TNF-α (**A**) and type-1 IFNs (**B**) across time in 10 stochastic replicates and their averages are plotted for both the chicken (blue) and duck (green) innate immune systems

Total IL-18 production in both chicken and duck AIIRABMs initially rapidly increased, then peaked, before subsequently decreasing (Fig.5). The chicken and duck average IL-18 trajectories were very similar: between the chicken and duck AIIRABMs peak IL-18 significantly differed by an average of 7 arbitrary units (one-way ANOVA, p < 0.05); chicken IL-18 did not peak at a significantly different time to duck IL-18 (one-way ANOVA, p = 0.22). The average duck IL-18 trajectory from 1 d.p.i. was always above the chicken trajectory, but it is difficult to determine whether the difference is biologically meaningful.

**Figure 5:**
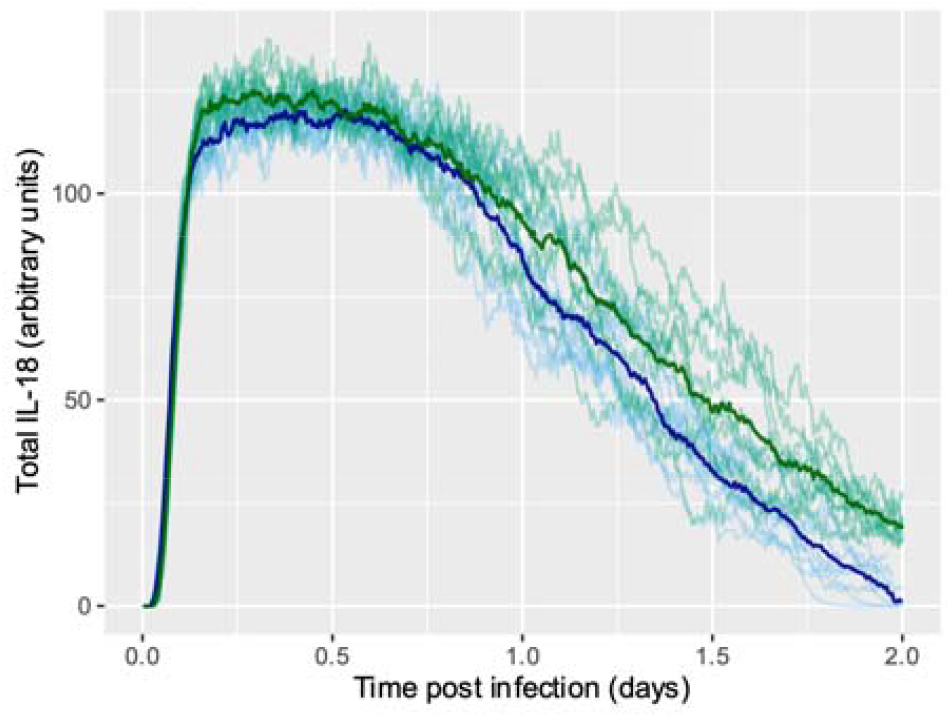
The AIIRABM calibrated, simulated across two days and at low initial inoculum (initial virus count = 50). Trajectories of total IL-18 across time in 10 stochastic replicates and their average are plotted for both the chicken (blue) and duck (green) innate immune systems.

### Parameter sweep of initial virus count

The parameter sweep of ‘initial virus count’ (representing HPAIV inoculation dose) was performed for an ‘initial virus count’ of 10 to 200 in the chicken and duck AIIRABMs. For each inoculation dose, the AIIRABM was simulated over two days with 10 stochastic replicates. In the chicken model total extracellular virus for all five inoculum doses initially increased, peaked, and subsequently decreased across the simulation period (Fig.6A). In the duck model the same relationship was observed for all inoculum doses except for ‘initial virus count’ 10 where extracellular virus never peaked (Fig.6B). In the chicken and duck systems extracellular virus peak intensity increased with inoculum dose (Kruskal-Wallis, p < 0.05; Kruskal-Wallis, p < 0.05). In both systems the time at which the extracellular virus peak occurred decreased with inoculum dose (Kruskal-Wallis, p < 0.05; Kruskal-Wallis, p < 0.05).

**Figure 6:**
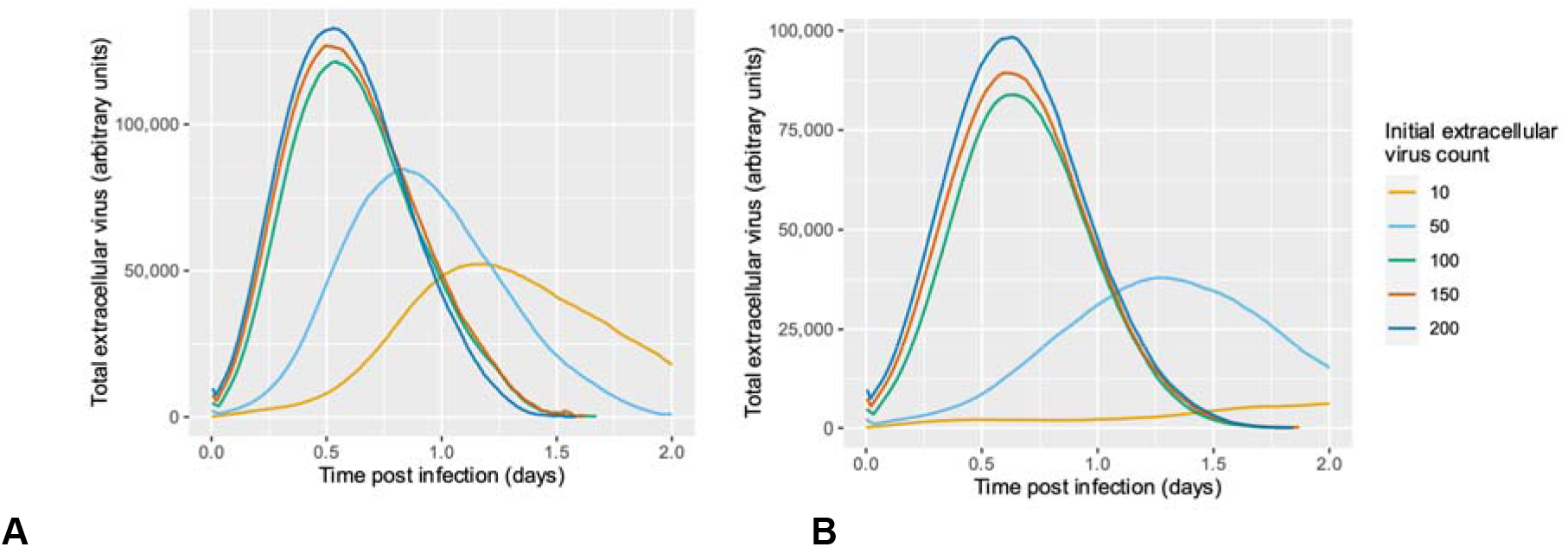
The AIIRABM calibrated, simulated across two days and parameter sweeping for inoculum dose (initial virus count from 10 to 200). Average trajectories of total extracellular virus across time in 10 stochastic replicates are plotted for both the chicken (**A**) and duck (**B**) innate immune systems.

In the chicken and duck systems there was limited to no significant difference between the peak intensities of inoculum doses 100, 150 and 200 (pairwise Wilcoxon rank sum tests, using p < 0.05 for statistical significance). Additionally, in both systems there was no significant difference between the times extracellular virus peaked for inoculum doses 100, 150 and 200 (Tukey test, using p < 0.05 for statistical significance). Therefore, increasing inoculum dose when the inoculum dose was already high had limited effect on infection severity and the time at which infection severity was greatest.

### *In silico* reinstatement of RIG-I into chickens

*In silico* reinstatement of RIG-I into chickens mimicked *in vivo* experiments with duRIG-I reinstatement into chickens via engineered expression vectors (preprint: Sid *et al*, 2023). The duRIG-I chicken AIIRABM was run across two days to mimic AIV *in vitro* cell culture experiments and simulated at low HPAIV inoculum dose with 10 stochastic replicates. The duRIG-I chicken AIIRABM stochastic replicates were compared to the chicken and duck AIIRABM stochastic replicates in section 3.1. The average trajectories for total extracellular virus, TNF-α and type-1 IFNs of the duRIG-I chicken AIIRABM closely followed the average trajectories of the duck AIIRABM (data not shown, see Fig.3 and Fig.4 for duck trajectories). The average IL-18 trajectory for the duRIG-I chicken AIIRABM more closely followed the duck average trajectory than the chicken trajectory by 1.5 d.p.i. (Fig.7). As mentioned in section 3.1, it is difficult to determine whether the difference in IL-18 production between the duck/duRIG-I chicken and chicken AIIRABMs is biologically meaningful.

**Figure 7:**
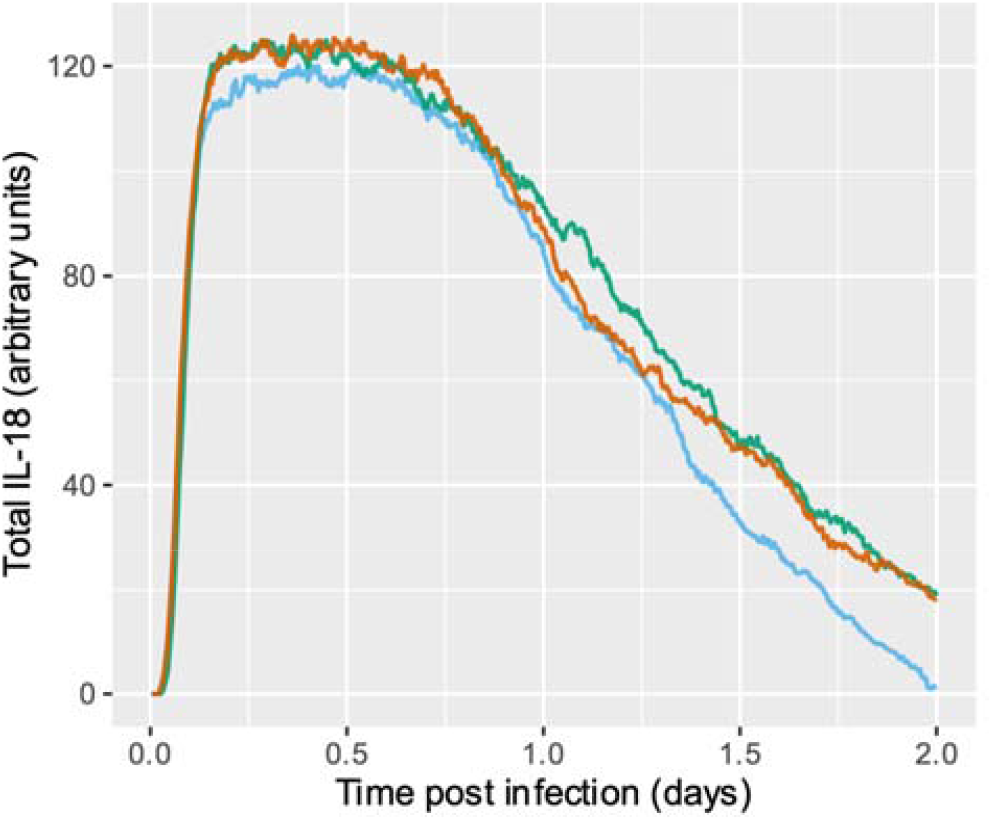
The AIIRABM with RIG-I reinstated into chickens, calibrated, simulated across two days and at low initial inoculum (initial extracellular virus count = 50). Average trajectories of total IL-18 across time in 10 stochastic replicates are plotted for the chicken (blue), duck (green) and duRIG-I chicken (brown) innate immune systems.

### The effect of the interaction between RIG-I and MDA5

The original AIIRABM assumed that RIG-I and MDA5 were functionally redundant in activating IRF7, and that RIG-I was a better IRF7 activator. The AIIRABM was altered to see whether the interaction type between RIG-I and MDA5 affected type-1 IFNs production. For all RIG-I MDA5 interaction types, type-1 IFNs production in both chicken and duck AIIRABMs increased to a peak before subsequently decreasing (Fig.8). In the original AIIRABM total type-1 IFNs peak intensity was greater in the duck AIIRABM than in the chicken (Kruskal-Wallis, p < 0.05; Fig.8A, duplicate of Fig.4B to enable easier comparison). Additionally, chicken type-1 IFNs peak production occurred earlier than duck peak production (Kruskal-Wallis, p < 0.05).

**Figure 8:**
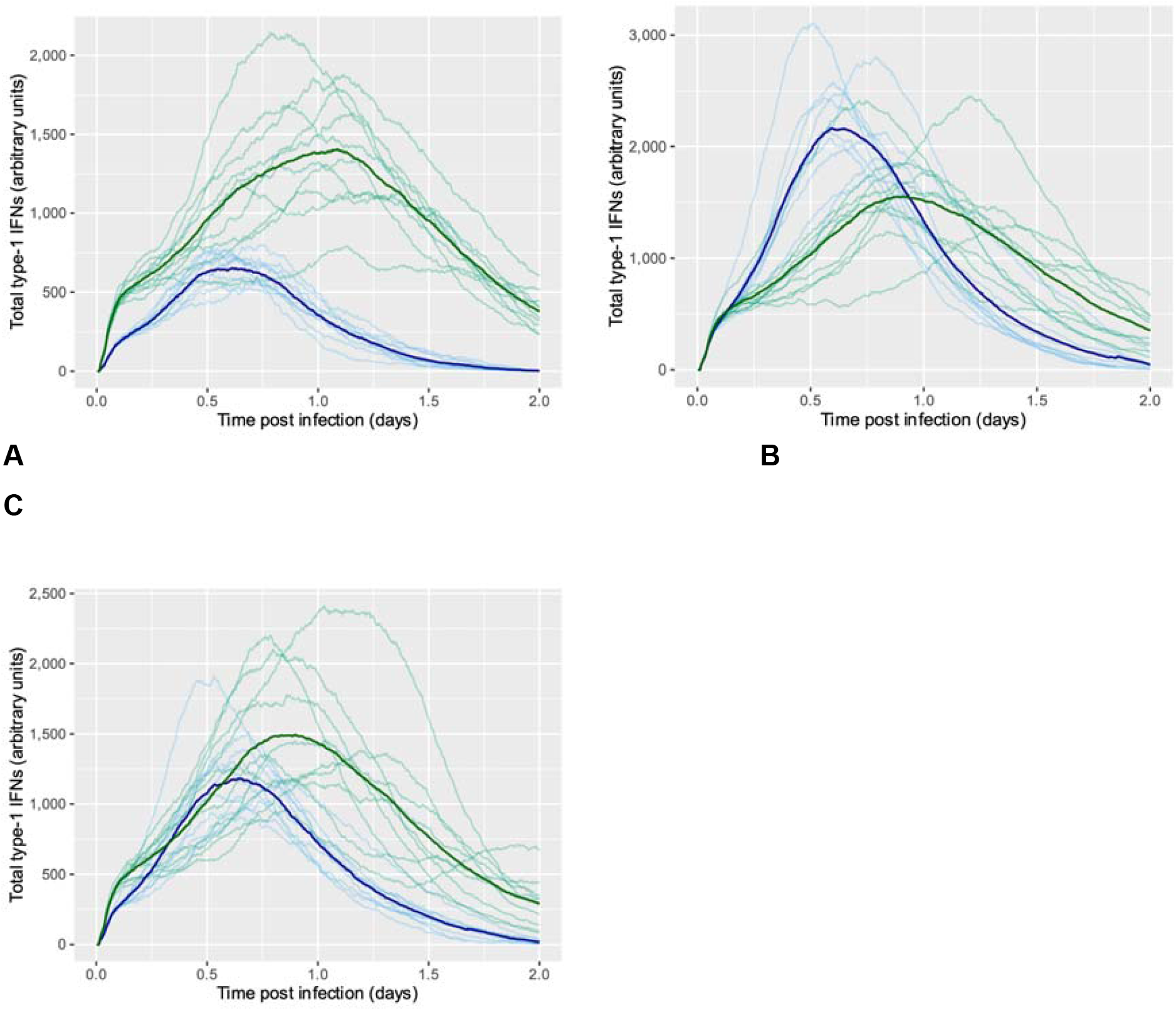
The three RIG-I MDA5 AIIRABMs calibrated, simulated across two days and at low initial inoculum (initial virus count = 50). Plots of the AIIRABM with: RIG-I and MDA5 functionally redundant and RIG-I a better activator of IRF7 (**A**); RIG-I and MDA5 functionally redundant and equal IRF7 activators (**B**); RIG-I and MDA5 equal IRF7 activators with a combined synergistic effect (**C**). Trajectories of total type-1 IFNs across time in 10 stochastic replicates and their average are plotted for both the chicken (blue) and duck (green) innate immune systems.

Two additional RIG-I MDA5 interactions were modelled: (1) functionally redundant, equally good at activating IRF7; (2) synergistic effect, equally good at activating IRF7. For the functionally redundant, equal activators AIIRABM, chicken type-1 IFNs production peaked much higher than duck production (one-way ANOVA, p < 0.05; Fig.8B). Additionally, chicken type-1 IFNs peak production occurred earlier than duck peak production (one-way ANOVA, p < 0.05). For the synergistic effect, equal activators AIIRABM, chicken and duck type-1 IFNs peak production intensity did not significantly differ (Kruskal-Wallis, p = 0.13; Fig.8C). Additionally, chicken type-1 IFNs peak production occurred earlier than duck peak production (one-way ANOVA, p < 0.05).

Between the three chicken AIIRABMs, peak type-1 IFNs production intensity significantly differed (Kruskal-Wallis, p < 0.05). However, the time at which production peaked did not significantly differ between the three chicken AIIRABMs (one-way ANOVA, p = 0.15). Between the three duck AIIRABMs, type-1 IFNs did not significantly differ in peal production intensity (one-way ANOVA, p = 0.49) or time of peak (one-way ANOVA, p = 0.76).

## Discussion

To our knowledge this is the first mechanism-based computational model that reconstructs the avian innate immune system response to HPAIV, and subsequently tests *in silico* the effects of key molecular differences in the chicken and duck immune systems. Both the chicken and duck AIIRABMs were successfully calibrated to mimic HPAIV *in vitro* cell culture experiments.

Increasing inoculum dose increases the probability of HPAIV infection in both chickens and ducks (Spekreijse *et al*, 2011; Swayne & Slemons, 2008). The same dose-response relationship was observed in the chicken and duck AIIRABMs where the maximum amount of extracellular virus produced increased with initial virus count. The greater virion production in chickens could increase the likelihood of macrophages and dendritic cells reaching phagocytic capacity (being overwhelmed with the number of virions to phagocytose) (Cline *et al*, 2017). Subsequently, the likelihood of an infectious particle migrating from the infection site to other tissues could increase, therefore increasing the likelihood of severe HPAIV infection in chickens. Increasing the inoculum dose within its highest range had limited effect on the infection severity and the time at which infection severity was greatest, suggesting beyond a certain inoculation dose infection severity is maximised. This phenomenon could be reflected *in vivo* by a mortality rate of nearly 100% beyond a given inoculum dose. A similar situation has been observed in chickens where the mean HPAIV infectious dose was equal to the mean lethal dose (Swayne & Slemons, 2008).

Differences in TNF-α production between the chicken and duck models agreed with experimental observations. TNF-α peaked in the chicken AIIRABM with an intensity nearly double that in the duck, agreeing with *in vitro* cell culture experiments where AIV induced greater pro-inflammatory cytokines production in chickens than ducks (Kuchipudi *et al*, 2014; de Bruin *et al*, 2022). The heightened TNF-α, pro-inflammatory response in chickens is thought to account for their severe HPAIV symptoms (Kuribayashi *et al*, 2013). Therefore, the AIIRABM further supports experimental observations of a heightened pro-inflammatory response in chickens contributing to greater HPAIV infection severity.

Whilst chickens generally show a pro-inflammatory response to HPAIV infection, ducks often produce an anti-viral state. Total type-1 IFNs peaked in the duck AIIRABM to a greater intensity and more rapidly than in the chicken AIIRABM. This result agrees with studies where chickens and ducks have been infected with HPAIV *in vivo* and ducks have produced greater quantities of type-1 IFNs (Cornelissen *et al*, 2013; El-Shall *et al*, 2023). Conversely other studies observed no significant difference in the type-1 IFN response between chickens and ducks (Kuchipudi *et al*, 2014) or that chickens produced a greater intensity of type-1 IFNs (Liang *et al*, 2011). However, the study that found chickens to be the greater type-1 IFNs producers also found that STAT-3 (an element of the JAK-STAT signalling pathway) was up-regulated in ducks with HPAIV infection and down-regulated in chickens. This could enable ducks to form a greater anti-viral state in their cells despite lower type-1 IFNs production. So, notwithstanding different experimental observations in the relative type-1 IFN responses in chickens and ducks with HPAIV infection, consensus suggests ducks successfully produce an anti-viral state which chickens cannot as easily achieve.

RIG-I is thought to be a key molecular difference that distinguishes the pro-inflammatory innate immune response of chickens and the anti-viral state of ducks with HPAIV infection (Karpala *et al*, 2011; Barber *et al*, 2010). Both extracellular virus and TNF-α production likely occurred at a lower rate in the duck AIIRABM than the chicken due to the inhibitory effect of RIG-I on virus replication. Type-1 IFNs, however, showed a more intense and rapid response in ducks despite type-1 IFNs being initially dependent upon viral nucleoproteins to stimulate their production, just like TNF-α. It is possible that the self-amplifying, positive feedback loop type-1 IFNs form (Erickson & Gale, 2008) enables them to become less dependent upon the levels of intracellular virus in the model, allowing their production to peak to a greater intensity in the duck AIIRABM than the chicken. It is interesting that when the interaction type between pattern recognition receptors RIG-I and MDA5 changed in the AIIRABM there was no effect on the duck type-1 IFNs production curve. This suggests that the presence alone of RIG-I in the duck model enables its greater type-1 IFN response. This hypothesis is further supported with the *in silico* reinstatement of duRIG-I into the chicken AIIRABM which increased chicken type-1 IFNs production so that it was similar to the production observed in the duck AIIRABM.

The AIIRABM results also suggest that RIG-I is a key molecular difference for affecting IL-18 production in chickens and ducks. Limited previous research on IL-18 production changes with HPAIV infection of fowl disagrees on whether chickens or ducks produce a greater IL-18 response (Kuchipudi *et al*, 2014; Tong *et al*, 2021). In the AIIRABM total IL-18 output by 2 d.p.i. was greater in ducks than chickens. The greater duck IL-18 response is likely only due to the presence of RIG-I in the duck AIIRABM as *in silico* reinstatement of duRIG-I in the chicken AIIRABM resulted in the IL-18 production more closely resembling the duck curve by 2 d.p.i. Microarray gene expression profiling of HPAIV infected chicken and duck macrophages could be conducted to experimentally validate the AIIRABM IL-18 response.

The baseline AIIRABM produced in this project agrees with previous work that RIG-I is seemingly of large importance to the high HPAIV immunity of ducks (Karpala *et al*, 2011; Barber *et al*, 2010). RIG-I transfection into the chicken AIIRABM as well as in previous experiments (Barber *et al*, 2010, 2013; preprint: Sid *et al*, 2023) has shown to improve the chicken immune response to HPAIV. *In silico* reinstatement of duRIG-I in the chicken AIIRABM resulted in outputs closely resembling those of the duck AIIRABM: the maximum extracellular virus count and pro-inflammatory response were lower in the duRIG-I chicken AIIRABM; type-1 IFNs production was greater in the duRIG-I chicken AIIRABM; IL-18 production was greater in the duRIG-I AIIRABM by the end of the *in silico* experiment, suggesting that Th1 cells would be greater stimulated and subsequently produce more IFN-γ. Note the different effect on the pro-inflammatory response with reinstating RIG-I in previous experiments and *in silico* with the AIIRABM. This discrepancy may result from the abstraction (simplification) of molecular processes in creating the AIIRABM, but provides an interesting area of research for further investigation requiring iterative rounds of experimental data collection and AIIRABM modelling.

However, the chicken type-1 IFN response changed with different RIG-I MDA5 interactions. When RIG-I and MDA5 were equally effective, redundant IRF7 activators, peak chicken type-1 IFNs production increased compared to the original chicken AIIRABM, peaking to a greater intensity than in the duck response. When RIG-I and MDA5 were modelled as equally effective activators, but synergistically functional (IRF7 is more likely to be activated if both pattern recognition receptors are active), peak chicken type-1 IFNs production did not significantly differ from duck type-1 IFNs production. These results suggest that the chicken type-1 IFN response is highly dependent upon the relative activation potential of MDA5, which is unsurprising considering the absence of RIG-I in the chicken system. Further experimental work needs to be done to confirm the difference in type-1 IFNs production between HPAIV-infected chickens and ducks (discussed above). If a consensus can be reached on whether HPAIV-infected chickens or ducks differ in their type-1 IFN responses, then the AIIRABM results could suggest the interaction type between RIG-I and MDA5. For example, if further research confirms ducks produce a greater type-1 IFN response, then the results from altering the AIIRABM would suggest RIG-I and MDA5 redundantly function to activate IRF7 and that RIG-I is the better activator.

The AIIRABM is limited in its representation of the avian inflammasome. Little is understood about the avian inflammasome and its signalling pathway, so the AIIRABM relies on our knowledge of the mammalian inflammasome to fill in any gaps. Whilst the mammalian inflammasome is relatively well understood, it differs significantly from the avian inflammasome. For example, many birds, including chickens and ducks, have lost the inflammasome ASC signalling molecule (the apoptosis-associated speck-like *protein* containing a caspase recruitment domain) (Billman *et al*, 2024). Therefore, the avian inflammasome, as in mammals, may be responsible for cleavage of pro-IL-18 into IL-18, but this interaction in birds has not yet been proven or its signalling pathway(s) determined. The AIIRABM assumes that an active avian inflammasome results in the cleavage of pro-IL-18.

Another limitation of the AIIRABM in its current form is that it models only the epithelial cells and macrophages of the avian innate immune response. Addition of other major cell types (i.e. Th1, dendritic and natural killer cells) to the AIIRABM could affect its molecular outputs. As well as adding in major cell types, the AIIRABM could improve by distinguishing M1 and M2 macrophage phenotypes. M1 and M2 macrophages result from different activating molecular signals (Yang *et al*, 2023). Currently the AIIRABM macrophages resemble M1 macrophages due to their direct involvement in the innate immune response. Including both M1 and M2 macrophages in the AIIRABM could provide further insight into the differences between the chicken and duck innate immune responses. However, despite its limitations, we believe that the AIIRABM provides a good initial approximation of the different effects of avian HPAIV infections between species and can provide a useful adjunct for future studies on comparative avian immunology, with possible extension to mammalian systems with a more direct potential impact on human health and future zoonotic pandemics.

## List of Abbreviations

AIIRABM: Avian Innate Immune Response Agent-Based Model
AIV: Avian Influenza Virus
d.p.i.: Days Post Infection
duRIG-I: Duck Retinoic Acid-Inducible Gene-I
HPAIV: Highly Pathogenic Avian Influenza Virus IFN Interferon
LPAIV: Low Pathogenic Avian Influenza Virus
RIG-I: Retinoic Acid-Inducible Gene-I

## Acknowledgements

This work was funded by UKRI BBSRC grants BB/Y007212/1 and BB/S001336/1 to BJF and CEB.

## Notes

### Competing Interest Statement

The authors have declared no competing interest.

